# Analyzing whole genome bisulfite sequencing data from highly divergent genotypes

**DOI:** 10.1101/076844

**Authors:** Phillip Wulfridge, Ben Langmead, Andrew P. Feinberg, Kasper D. Hansen

## Abstract

In the study of DNA methylation, genetic variation between species, strains, or individuals can result in CpG sites that are exclusive to a subset of samples, and insertions and deletions can rearrange the spatial distribution of CpGs. How to account for this variation in an analysis of the interplay between sequence variation and DNA methylation is not well understood, especially when the number of CpG differences between samples is large. Here we use whole-genome bisulfite sequencing data on two highly divergent inbred mouse strains to study this problem. We find that while the large number of strain-specific CpGs necessitates considerations regarding the reference genomes used during alignment, properties such as CpG density are surprisingly conserved across the genome. We introduce a method for including strain-specific CpGs in differential analysis, and show that accounting for strain-specific CpGs increases the power to find differentially methylated regions between the strains. Our method uses smoothing to impute methylation levels at strain-specific sites, thereby allowing strain-specific CpGs to contribute to the analysis, and also allowing us to account for differences in the spatial occurrences of CpGs. Our results have implications for analysis of genetic variation and DNA methylation using bisulfite-converted DNA.

## Introduction

DNA methylation is a key epigenetic mark that has become widely implicated in human development and disease [1, 2]. Accurate determination of methylation at CpG dinucleotide positions across the genome is critical for understanding its association with functional regulation. Multiple techniques currently exist to perform this measurement, each with varying degrees of genomic coverage and depth. One gold-standard method is whole-genome bisulfite sequencing (WGBS), which pairs bisulfite conversion of cytosine residues with next-generation sequencing [3]. To quantify methylation in WGBS, bisulfite converted reads are compared to an in silico bisulfite-converted genome sequence, referred to as the reference genome [3, 4, 5]. At each CpG site in the reference genome, an aligned read is called as unmethylated if the sequence is TG (indicating bisulfite conversion) and methylated if the sequence is CG (indicating protection by the methyl group). Statistical packages such as BSmooth [4] can then integrate this data across larger regions to estimate and compare overall methylation patterns between sample groups.

It is well understood that a CG-to-TG mutation is the most common dinucleotide mutation in the mammalian genome, due to the high rate of spontaneous deamination at methylated CpGs [6, 7, 8]. This effect becomes particularly pronounced in animal models such as mice: inbred strains can be separated by up to millions of nucleotide variants and indels, a large proportion of which affect CpG dinucleotides. This presents a substantial pitfall when analyzing bisulfite converted DNA: if the sample genome has a CG-to-TG (or CG-to-CA) variant relative to the reference genome sequence, reads aligning to the variant using standard alignment approaches will produce unmethylated calls without inducing any alignment mismatches [9]. This will result in an excess of 0% methylated “sites” at locations where there is actually no CpG sequence.

The problematic effect of C/T variants on bisulfite alignment and methylation quantification is widely recognized, and various methods exist to compensate. While it is possible to identify variants in bisulfite converted data by considering reads aligned to the opposite strand [9, 10], it is self-evident that the best solution is to simply align each sample to its own genome sequence, assuming such sequence is available. Variants of this strategy are indeed commonly used in analysis, i.e. with alignment to approximate personal genomes; this has been done both in plants [11, 12, 13] and mammals [14, 15]. However, while both the problem and the optimal solution are recognized, the magnitude of the bias is not well understood or described. Filling this gap in understanding is especially important for experiments involving a large number of distant or mixed genotypes. In these experiments, knowing whether CpG variation presents a substantial enough bias to necessitate the computational cost of acquiring tens to hundreds of personal genomes, which can demand substantial resources, is of high importance.

But the effect of C/T variants goes beyond alignment. Specifically, if each sample is aligned to a separate genome, a critical issue arises on how comparisons should be made across different genomes, whose coordinates will be separated by insertions and deletions, as well as how strain- or sample-specific CpGs (that is, CpGs only existent in a subset of individuals) should be handled in analysis. Common methodologies treat sample-specific CpGs as incomparable and either drop CpGs not covered in all samples, or drop samples for which a CpG does not exist. As a recent example, a study in humans [14] produced approximate sample-specific genomes by substituting single nucleotide variants into a common reference genome, then analyzed methylation differences at individual CpGs; for each of these CpGs, samples in which the CpG was lost were discarded from analysis. In an analysis that drops sample-specific CpGs, however, a genomic location where half the samples had a CpG with high methylation, while the other half had lost the CpG, would not be identified as a position of variable methylation.

In the face of mounting evidence that sites of hypervariable CpG mutation are strongly related to species-specific disease and phenotype [16], failing to account for the functional implications of CpG loss may represent a substantial loss of information. It seems clear that an approach that fully factors in the presence of sample-specific CpGs will produce a more complete analysis. But it is currently an open problem how such an analysis should be performed.

Here, we address two basic questions. First, how essential is it to use sample specific genomes, since acquiring such data comes at a cost? Second, how should different coordinate systems across reference genomes be reconciled, and how can strain- or sample-specific CpGs be factored into comparisons? Specifically, we examine sequence variation and methylation data from two inbred but highly divergent mouse strains: C57BL/6J, upon which the mouse reference genome is based, and the wild-derived CAST/EiJ. We describe patterns in strain-specific CpG variation, and quantify the effect of ignoring this variation when performing alignment and methylation quantification. Next, we propose a method that incorporates strain-unique CpGs into comparative analysis, allowing them to contribute to identification of differentially methylated regions, and show that doing so increases power. Our method is based on using smoothing to impute methylation levels at strain-specific CpGs.

## Results

### WGBS data

As an extreme example of massive CpG variation between genomes, we examined sequence and methylation data from two inbred mouse strains. C57BL/6J (BL6) is the standard laboratory strain and the basis of (i.e. equivalent to) the mm9 reference genome, whereas CAST/EiJ (CAST) is wild-derived and highly divergent, as evidenced by a large number of single nucleotide variants (20,539,633) and indels (8,171,218), and has a reference genome available for use in alignment [17]. Methylation data was generated via WGBS on liver samples from each strain (n = 4 per strain). The initial purpose of the sequencing experiment was to identify strain-specific regions of differential methylation, possibly driven by genetic changes. We generated sequencing data at relatively low coverage (Table 1) and aligned all samples to both genomes using Bismark (Methods).

**Table 1.** Number of reads and alignment statistics.

| Sample | nReads | BL6 |  |  | CAST |  |  |
| --- | --- | --- | --- | --- | --- | --- | --- |
|  |  | nAligned | aRate<sup>a</sup> | CpG<sup>b</sup> | nAligned | aRate<sup>a</sup> | CpG<sup>b</sup> |
| BL6_1 | 165,455,305 | 92,198,446 | 55.7 | 5.8 | 81,398,771 | 49.2 | 4.7 |
| BL6_2 | 155,446,345 | 85,632,105 | 55.1 | 5.5 | 75,566,778 | 48.6 | 4.5 |
| BL6_3 | 165,687,191 | 99,880,547 | 60.3 | 6.8 | 87,713,251 | 52.9 | 5.5 |
| BL6_4 | 172,926,402 | 101,892,159 | 58.9 | 6.7 | 89,939,416 | 52.0 | 5.5 |
| CAST_1 | 171,357,014 | 107,621,399 | 62.8 | 6.3 | 121,678,485 | 71.0 | 6.4 |
| CAST_2 | 161,768,892 | 90,231,861 | 55.8 | 6.1 | 102,071,949 | 63.1 | 6.3 |
| CAST_3 | 154,973,730 | 86,822,723 | 56.0 | 5.5 | 98,305,382 | 63.4 | 5.7 |
| CAST_4 | 188,134,260 | 90,420,870 | 48.1 | 6.0 | 102,599,995 | 54.5 | 6.2 |

<sup>a</sup>alignment rate
<sup>b</sup>coverage of CpGs

### CpG variation across the mouse genome

We first examined differences between the DNA sequences of BL6 and CAST in order to quantify and characterize CpG variation. To facilitate accurate sequence comparisons between these two genomes, which do not share coordinate systems, we used the modmap tool [18]. This tool functions similarly to UCSC’s liftOver [19] to convert a set of genomic locations to their corresponding coordinates in another strain. Using this tool, we can determine whether a CpG in one strain retains its sequence in the other strain, contains a mutation, or cannot be accurately mapped, which occurs when an indel over a CpG results in an ambiguous position after modmap conversion (Figure 1a).

**Figure 1.**
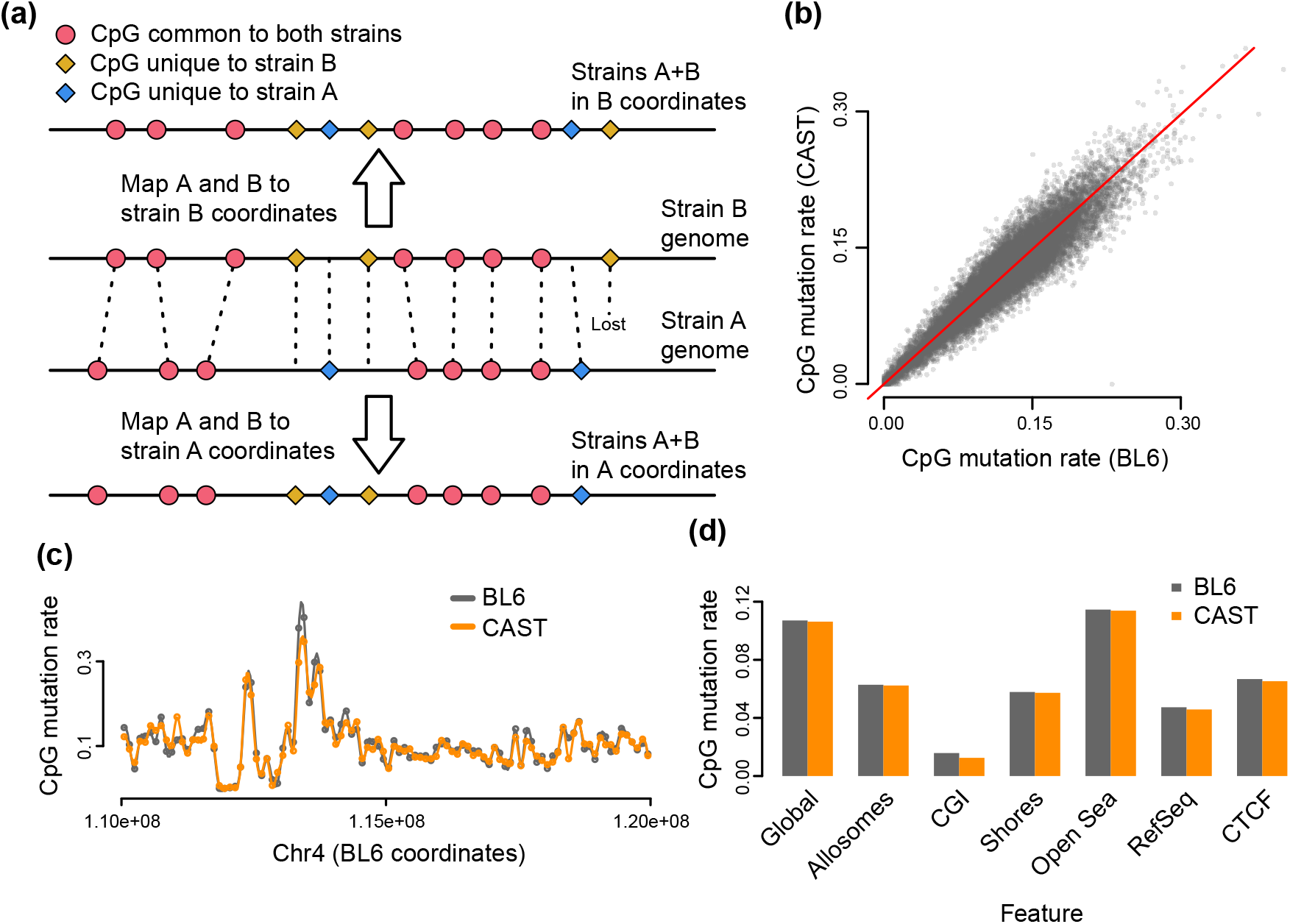
Characteristics of strain-specific CpG mutations. (**a**) A representation of whole-genome alignment between two strains, A and B. The CpGs present in the two strains can be represented in both coordinate systems; CpGs are either shared between strains (red), mutated (yellow or blue), or “lost” due to indels that render the coordinate unmappable (rightmost yellow). (**b,c**) The CpG mutation rate (proportion of strain-specific CpGs relative to total CpGs) calculated in 100kb bins, is comparable between BL6 and CAST strains, both genome-wide (**b**) and locally (**c**). (**d**) The CpG mutation rate across different genomic features.

Using modmap and the FASTA files for mm9 (BL6) and CAST, we extracted and tabulated the forward-strand dinucleotide sequences corresponding to the 21.3 million CG dinucleotides in BL6; the results are shown in Table 2. Approximately 19 million CGs from BL6 are shared by CAST; 1.64 million are mutated to either TG or CA, with another 0.54 million mutated to GG/CC and AG/CT. Another 100k CpGs could not be accurately mapped from BL6 to CAST due to strain indels over the sequence. We observed similar results when we performed the reverse analysis, tabulating sequences of CAST CpGs when modmapped to BL6: of 21.4 million CGs in CAST, about 2.3 million are unique to CAST, with a differing or unmappable sequence in BL6 (Table 2).

**Table 2.** Number of CpGs in different strains.

|  | Autosomes |  | Allosomes |  |
| --- | --- | --- | --- | --- |
|  | BL6 | CAST | BL6 | CAST |
| Common CG | 18,140,628 |  | 916,392 |  |
| TG/CA in other strain | 1,601,446 | 1,669,901 | 43,330 | 45,041 |
| Lost in other strain | 522,845 | 535,219 | 15,394 | 15,838 |
| Unmappable | 99,781 | 90,356 | 2,676 | 2,438 |
| Total | 20,364,700 | 20,436,104 | 977,792 | 979,709 |

Based on our observations, we calculated the ratio of strain-unique to total CpGs, or CpG “mutation rate,” at roughly 2.3M/21.3M=10.7%, which can be compared to the overall genomic mutation rate of roughly 1.2% (computed using variant and indel information from modmap; see Methods). It is well understood that this high rate of CpG mutation is caused by the nature of DNA methylation, as methylated C positions are more likely to undergo spontaneous deamination to T [7, 8, 6]. The CpG mutation rate is not uniform throughout the genome, but can fluctuate across local regions. Interestingly, however, the proportion of BL6-unique CpGs in any given region tends to closely match that of CAST-unique CpGs (Figure 1b, c). The CpG mutation rate also varies between different functional regions, for example being much lower in CpG islands and somewhat lower in promoter and CTCF sites (Figure 1d).

### Not using strain specific genomes induces dramatic mapping bias

To show the impact of not using strain specific genomes for alignment, we first aligned sequencing reads from both strains to the standard BL6/mm9 reference genome, which represents an analysis pipeline making no adjustment for genotype differences. When we computed the average methylation across all read-covered autosomal CpGs in the reference, which we call global methylation, the two strains showed a large difference, with the CAST strain’s estimates lower by over 7.6% (Figure 2a, p < 1.3 × 10 ^−6^). This difference is comparable in magnitude to the level previously observed between tumor and normal colon [20] and associated with EBV-mediated oncogenesis [21], and is far larger than what we could realistically expect from a comparison between strains or individuals. Note that global methylation is an average across millions of CpGs, and thus unlikely to be affected to this extent by local differences in coverage. This observation is reversed when all samples are aligned to the CAST genome (Figure 2a), which proves this is caused by the choice of genome used for alignment and shows that the bias introduced by using the wrong genome is 7-8%.

**Figure 2.**
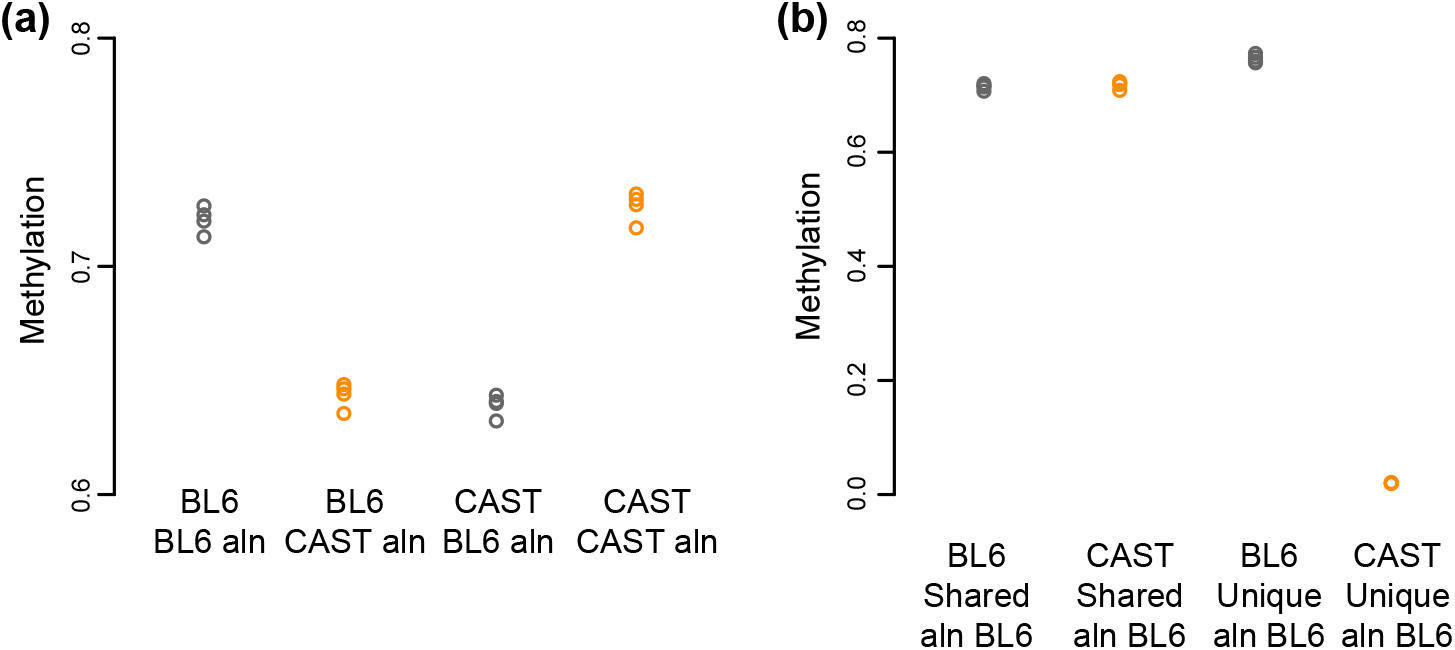
Effect of CpG variation on global methylation. (**a**) Global methylation estimated for four samples each of two different mouse strains (BL6 and CAST) aligned to either of the two reference genomes. Samples aligned to the strain-specific genome display higher global methylation compared to the same sample aligned to a distant reference genome. (**b**) Average methylation for different sets of CpGs for different samples, all aligned to the same reference genome. CpGs only present in CAST are quantified as having 0% methylation when aligned to BL6.

As expected, the observed discrepancy in global methylation can be explained by CpG sequence variation. When we categorized CpGs by whether they were present in both strains, we observed a large methylation difference only in the CpGs unique to BL6, with the measured methylation levels of CAST samples in those CpGs being virtually 0% (Figure 2b). Unsurprisingly, this is addressed by alignment to personalized reference genomes: when reads from the CAST samples were aligned to the CAST reference genome, their global methylation was roughly equivalent to those of BL6 samples aligned to mm9 (Figure 2a). This shows that the bias in global methylation is predominantly caused by assigning a methylation percentage of 0% to a “lost” CpG.

As with global methylation, analysis of focal changes can also be strongly impacted by the presence of CpG variation combined with mapping bias. As an example of focal analysis, we used the BSmooth pipeline [4] to identify small differentially methylated regions (DMRs) between BL6 and CAST, which range from hundreds to a few thousand bases in width. Performing this analysis on mm9-aligned samples, without adjustment for CpG variation, identified 2,865 DMRs that passed our cutoff criteria, which includes a measure of family-wise error rate (gFWER ≤ 1/18, mean difference > 0.1; see Methods).

However, the vast majority of these DMRs are artifacts of misleading 0% methylation measurements at BL6-unique CpGs in CAST samples. When we simply removed these BL6-unique CpGs prior to smoothing as a rudimentary form of adjustment, many observed mean differences in DMRs were diminished or absent entirely - only 554 (19.3%) were still identified as significant by the new analysis (Figure 3c). As further evidence of the impact of mapping bias, many DMRs reversed in direction when we aligned all samples to the CAST genome and performed focal analysis, as BL6 samples now had a disproportionate number of 0% methylation measurements (Figure 3b). In all of these cases, the magnitude of a given DMR’s methylation discrepancy was highly correlated with the number of unique CpGs in the region (Figure 3d). This analysis shows that mapping bias can caused false positives which vastly outnumbers the potential true positive strain differences. It also suggests the presence of a modest number of genomic regions which are differentially methylated between the two strains – the original purpose of generating the data.

**Figure 3.**
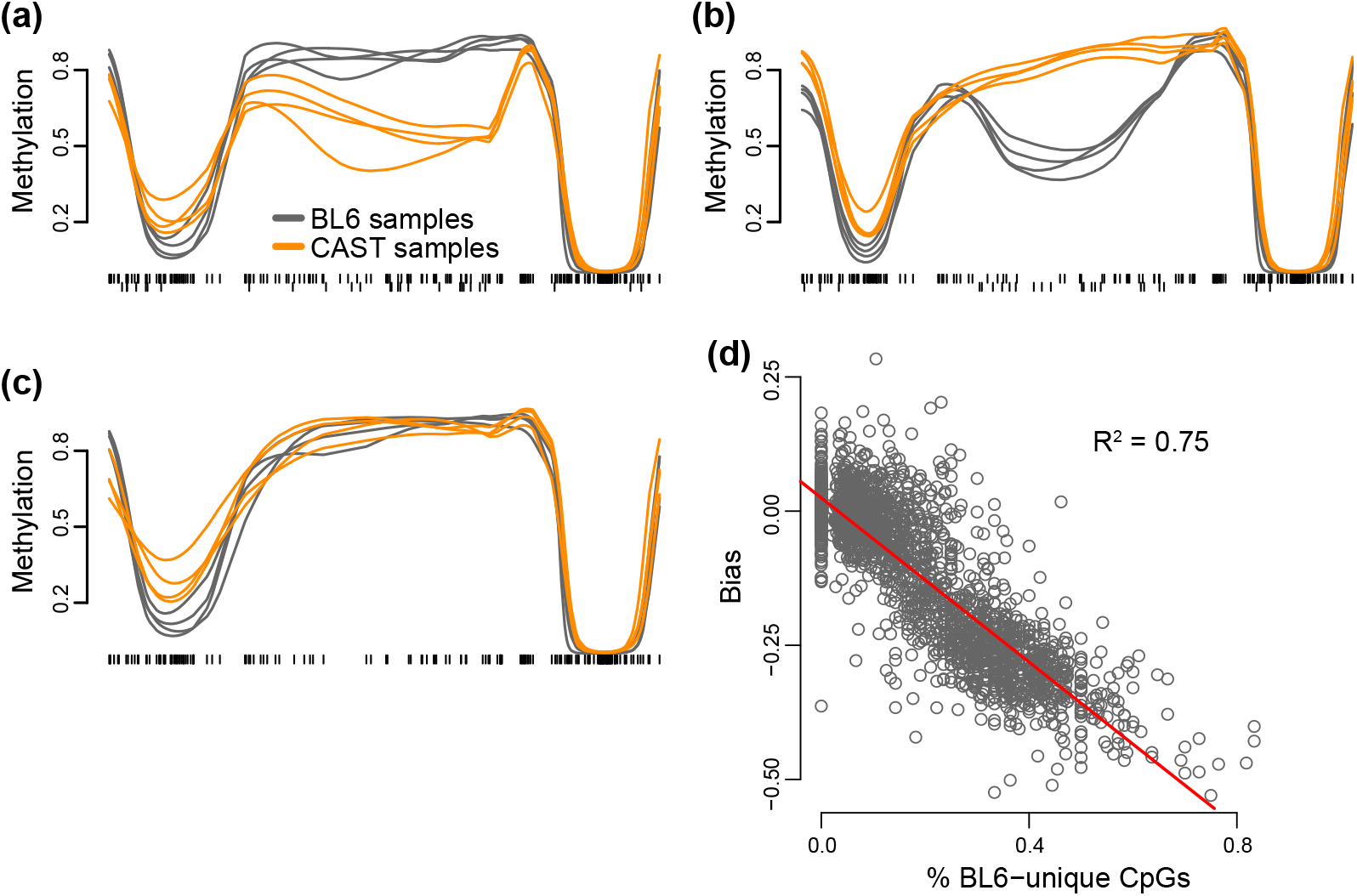
Mapping bias causes false-positive focal methylation changes. The example region pictured in (**a**) is identified as a DMR in mm9-aligned analysis, due to multiple BL6-unique CpGs (downwards ticks) that are read as having near-0% methylation in CAST samples. The DMR is reversed in direction if all samples are aligned to CAST due to the same effect occurring with CAST-unique CpGs (**b**), and no differential methylation is observable once these sites are removed from analysis (**c**). The magnitude of bias in such false-positive DMRs (i.e. the change in apparent differential methylation when unique CpGs are removed) is highly correlated with the regional level of CpG variation (c).

We have previously described an analysis method for identifying large-scale changes in DNA methylation, at the scale of 100kb or more CITE. Apply this method to data aligned to BL6 identified 2,354 regions comprising 347 Mb and covering 1.9M CpGs passing our cutoff criteria. For data aligned to CAST we found 4,199 regions comprising 1,101 Mb and 6.4M CpGs. For this analysis, the direction of change was fully consistent with the direction implied by differences in global methylation: 98.6% of the regions were hyper-methylated in BL6 when aligned to BL6 and 99.3% were hypermethylated in CAST when aligned to CAST. Figure 4 illustrates a reversible large-scale change. Unlike the result for small scale changes, this analysis suggests that there are no large-scale changes in DNA methylation between the two strains.

**Figure 4.**
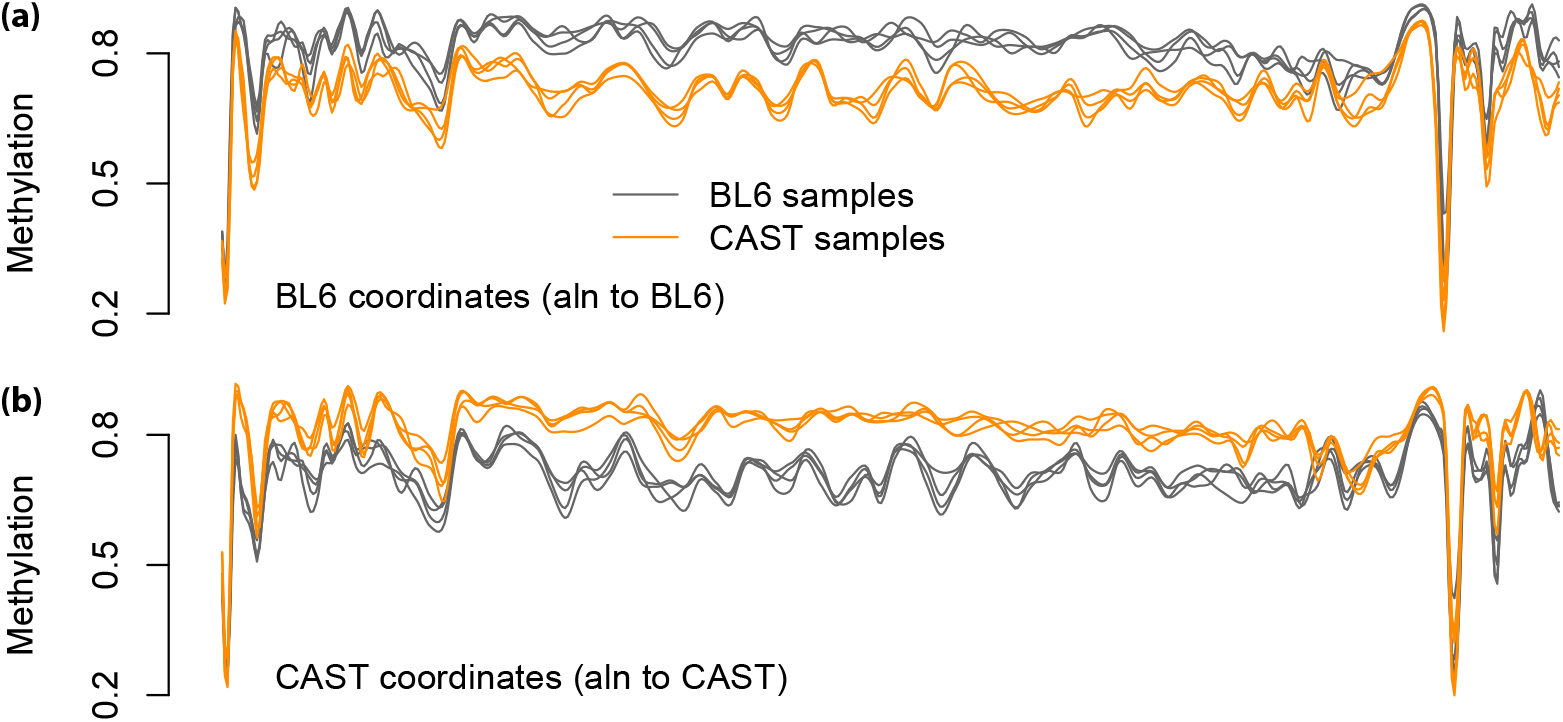
Mapping bias causes false-positive large-scale methylation changes. The same 2.4 Mb genomic region is depicted in two different coordinate systems and with different data processing. (**a**) All samples aligned to BL6. (**b**) All samples aligned to CAST.

Based on these observations, we conclude that use of personalized reference genomes, or at least careful consideration of adjustments for CpG variation, is imperative when performing WGBS analysis on samples with distant genotypes.

### Regional smoothing over strain-unique CpGs increases power for focal analysis

We now discuss potential strategies for dealing with sites of CpG variation. Essentially, two options exist: remove all CpG sites only present in one genotype, or somehow include those sites indirectly. The former strategy is straightforward to implement and widely used [22], but (as we have noted above) sacrifices a substantial amount of potential information. In contrast, the latter is only possible for regional rather than single-position analyses, and even then some method must be designed to allow indirect comparison of nearby (but offset) genotype-unique CpGs.

Fortunately, existing methods such as imputation via smoothing, which are already designed for regional analysis, are primed to handle data from genotype-unique CpGs. For example, BSmooth [4] can smooth methylation data over all CpGs without a need to manually designate genotype-unique CpGs: under default settings, CpG sites that are nonexistent in a certain sample are simply treated as zero-coverage, and thus are excluded from smoothing in that sample. The BSmooth process yields a set of continuous methylation curves across the whole genome, which can then be evaluated within just a set of common CpGs; however, data from nearby genotype-unique CpGs influence the imputed methylation values at these common CpGs, and thus still contribute to the final analysis.

Specifically, we propose the following algorithm:

1. Map reads to individual (strain-specific) genomes, and quantify methylation.
2. Using whole-genome alignment tools, such as modmap or liftOver, map CpGs between personalized genomes, into a common coordinate space.
3. Smooth methylation values to obtain personalized, continuous methylation profiles in the coordinate system of choice.
4. Analyze the methylation profiles to obtain differentially methylated regions.

Using the same WGBS data from BL6 and CAST mice as above, we tested what we will refer to as “unique-removed” and “unique-included” analysis pipelines. As before, we aligned BL6 samples to the BL6/mm9 reference genome and CAST samples to the CAST reference genome, then used modmap to place all samples in the mm9 coordinate system. For the unique-removed analysis, we removed 4.4 million CpGs unique to either BL6 or CAST, then smoothed over the remaining 19 million CpGs using BSmooth. In contrast, all 23.4 million CpGs were retained and smoothed in the unique-included analysis, as per the algorithm outlined above. Focal analysis was then performed to identify small DMRs.

Strikingly, more differential methylation is identified when strain-unique CpGs are included in analysis. Unique-included analysis found 976 DMRs passing our cutoff criteria (gFWER ≤ 1/18, mean difference > 0.1; Methods), while unique-removed analysis found 716 DMRs. There was no complete overlap between these two groups; 321 DMRs in the unique-included analysis were not present in unique-removed analysis, while 61 unique-removed DMRs were not identified by unique-included analysis. As might be expected, these differences appear to result from the additional effect that strain-unique CpGs exert on imputed methylation: while DMRs identified by both analyses tend to agree between analyses in terms of the mean difference in smoothed methylation of those regions, DMRs identified by only one analysis almost universally have a higher computed mean difference in that analysis (Figure 5a, b).

**Figure 5.**
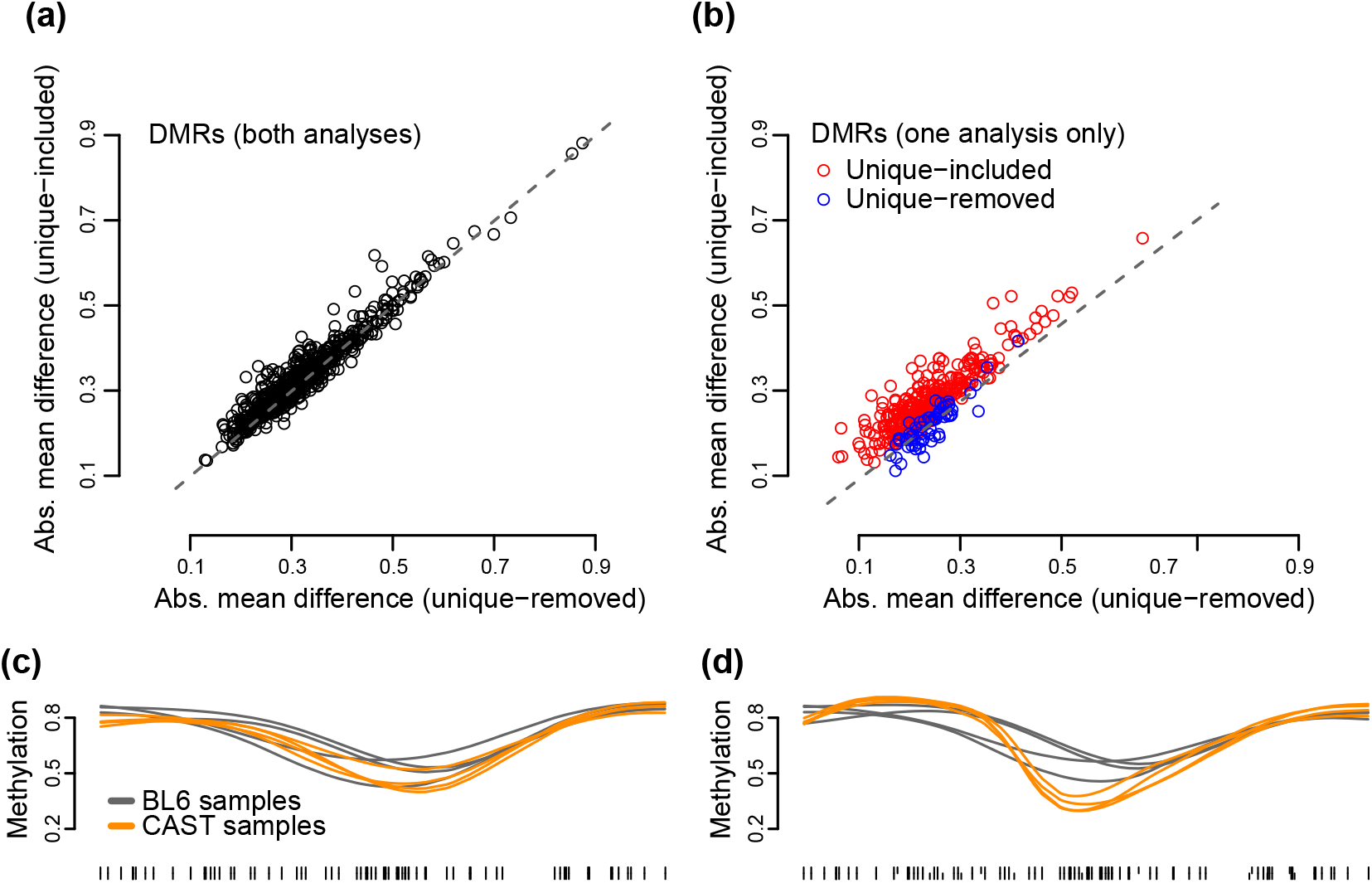
Including strain-unique CpGs identifies additional DMRs in focal analysis. DMRs identified in both unique-removed (shared CpGs only) and unique-included (all CpGs, including strain-unique) strategies show roughly similar mean differences in smooth methylation between strains, when computed by either analysis (**a**). In contrast, regions identified in only one analysis show a much higher mean difference in that analysis (**b**). An example DMR chosen from those only identified in the unique-included analysis (b, red) shows no observable difference between strains in the unique-removed analysis (**c**) but a visible difference in the unique-included analysis (**d**).

As an example of a DMR only identified by unique-included analysis, we plotted a region within the 5730522E02Rik gene, which has been previously implicated in mouse strain-specific expression [23]. When smoothing only over shared CpGs in the unique-removed analysis, there is no observable difference between BL6 and CAST in this region (Figure 5c). However, when strain-unique CpGs are kept in analysis, the resulting smoothing annotates a 21% methylation difference between strains (Figure 5d). Querying the same region in unique-removed analysis yields a 6% methylation difference which is below detection level. We conclude that including strain-specific CpGs in the analysis increases power.

To further characterize regions identified by each analysis, we examined enrichment of overlaps between DMRs and various functional marks obtained from ENCODE, as well as genomic features of interest such as Refseq gene promoters and CpG islands. The results, in log2 odds-ratios, are summarized in Table 3. All sets of DMRs, whether identified in both analyses or unique to one, had similar levels of enrichment for H3K4me1, H3K4me3, and H3K27ac (histone modifications enriched at enhancers and promoters), as well as for CTCF (a methylation-sensitive transcription factor). Strikingly, DMRs identified only in the unique-included analysis were uniquely depleted for Pol2 and CpG islands, suggesting that unique CpGs contribute to differential methylation away from promoter regions; this is consistent with the observation that the mutation rate is low at CpG islands (Figure 1d). Overall, our enrichment analysis indicates that the DMRs identified only by the unique-included analysis are functionally relevant in liver, and represent true additional regions of interest. The enrichment in DMRs identified only in unique-removed analysis is slightly more difficult to interpret; we suggest that these could represent regions that, although they may be in functional areas, drops below detection level when strain-unique CpGs are removed.

**Table 3.** Enrichment of strain-specific DMRs in genomic features.

| Feature | log2(OR) |  |  | *p*-value |  |  |
| --- | --- | --- | --- | --- | --- | --- |
|  | Shared | Included only | Removed only | Shared | Included only | Removed only |
| Refseq promoters | 0.98 | 0.55 | 0.23 | $< 2.2 \cdot 10^{-16}$ | $< 2.2 \cdot 10^{-16}$ | 0.046 |
| CpG Islands | 0.52 | -0.42 | 0.099 | $< 2.2 \cdot 10^{-16}$ | $4.76 \cdot 10^{-5}$ | 0.59 |
| CpG Shores | 1.21 | 1.09 | -1.52 | $< 2.2 \cdot 10^{-16}$ | $< 2.2 \cdot 10^{-16}$ | $3.4 \cdot 10^{-10}$ |
| H3K4me1 | 3.11 | 2.92 | 3.81 | $< 2.2 \cdot 10^{-16}$ | $< 2.2 \cdot 10^{-16}$ | $< 2.2 \cdot 10^{-16}$ |
| H3K4me3 | 1.99 | 1.48 | 2.16 | $< 2.2 \cdot 10^{-16}$ | $< 2.2 \cdot 10^{-16}$ | $< 2.2 \cdot 10^{-16}$ |
| H3K27ac | 1.97 | 1.96 | 2.60 | $< 2.2 \cdot 10^{-16}$ | $< 2.2 \cdot 10^{-16}$ | $< 2.2 \cdot 10^{-16}$ |
| CTCF | 1.59 | 1.74 | 2.14 | $< 2.2 \cdot 10^{-16}$ | $< 2.2 \cdot 10^{-16}$ | $< 2.2 \cdot 10^{-16}$ |
| Pol2 | 0.32 | -1.28 | 1.33 | $1.2 \cdot 10^{-7}$ | $< 2.2 \cdot 10^{-16}$ | $1.5 \cdot 10^{-15}$ |
| Any feature above | 3.09 | 2.87 | 3.54 | $< 2.2 \cdot 10^{-16}$ | $< 2.2 \cdot 10^{-16}$ | $< 2.2 \cdot 10^{-16}$ |

### Strategies for methylation analysis in the absence of personal genomes

In the previous sections, we have discussed differential analysis of divergent genomes under the assumption that accurate reference genomes exist for each unique genotype. Though we expect this to be the case for most inbred mouse strains, which are extensively curated, we can envision situations (e.g. studies in other species) where such personal references are unavailable; in these cases, other measures must be taken to mitigate alignment bias.

One possibility for addressing CpG variation is genotyping of samples using bisulfite converted data, for which various methods exist [9, 24, 10]. However, these methods have been evaluated on human data, where the number of SNVs in an individual is roughly an order of magnitude smaller than that between the more distant mouse strains - 3.3 million SNPs in human [25], compared to 20.5 million SNPs between BL6 and CAST (Methods). Furthermore, these methods require high coverage to work. Nevertheless, we proceeded to assess this approach for our data.

To test the feasibility of bisulfite genotyping in our data, we pooled our 4 genetically identical CAST replicates into one meta-sample with 24x coverage, aligned to the BL6 genome, and genotyped using BS-SNPer [10]; we then examined how much of CAST’s CpG variation was accurately identified. From our CAST alignment data, we know that CAST-unique CpGs in the meta-sample are well-covered by reads, with 2.17M of 2.26M covered at the recommended 10x or higher; despite this, however, BS-SNPer performed poorly at identifying these CpGs, only calling 1.15M (51%) correctly (Table 4). Identification of BL6-unique CpGs, which are lost in the CAST metasample, was even less efficient, with only 542K of 2.29M (24%) identified by BS-SNPer as non-CpG; this is perhaps expected, given that in a CG-to-TG mutation half of the reads are uninformative for genotyping under bisulfite conditions. Accordingly, even after adding and removing CpGs called by BS-SNPer, our calculated global methylation for the CAST metasample of 68.2% remained markedly lower than those obtained from personal alignments (72-73%), indicating mapping bias was incompletely addressed. Together, our results suggests that bisulfite genotyping was insufficient for addressing CpG variation between our divergent strains, at our sequencing depth.

**Table 4.** BS-SNP results on pooled CAST samples.

|  |  |
| --- | --- |
| BL6-unique CpGs | 2,285,477 |
| BL6-unique CpGs identified by BS-SNPer | 542,642 |
| CAST-unique CpGs | 2,266,009 |
| CAST-unique CpGs identified by BS-SNPer | 1,153,611 |

Another potential approach to dealing with CpG differences is to align to the standard reference, then filter out known variation based on an external database, such as dbSNP. For example, one could exclude any CpG overlapping known SNPs of certain criteria (high population frequency, CG-to-TG, etc.). Assuming a sample has a relatively small number of private SNPs, this method would theoretically remove the main source of hypomethylation bias, and indeed this method has been previously used in literature, albeit on much less distant genotypes [26]. However, stringent filtering could overestimate the amount of variation and remove a proportion of CpGs that remain in the sample, in addition to truly lost CpGs. Furthermore, non-CpG differences from the reference would not be addressed by this method.

To test this approach, we aligned CAST samples to the BL6 reference genome, then removed all BL6-unique CpGs from the dataset prior to smoothing; this represents a “perfect” adjustment, in which all CpG variation is removed with no overestimation. As expected, this method removed the bias in global methylation, with estimates close to those obtained with personal genomes (Figure 6a). Focal analysis identified 622 DMRs, of which 529 overlapped with the 716 DMRs identified in the personal, unique-removed analysis, indicating that differential finding under this method is at least somewhat comparable, though with some loss of power. However, care must still be taken when examining results obtained with this approach, as non-CpG-related alignment errors can still produce false-positive regions. As an example, we found a region on chr10 containing no strain-unique CpGs, which appeared to be greatly hypomethylated in CAST samples when aligned to the BL6 reference, but showed no methylation difference when samples were aligned to personal references (Figure 6b). We discovered that this result was due to thousands of unmethylated CAST mitochondrial reads, which aligned preferentially to the BL6 chr10 rather than to the BL6 chrM due to SNPs in both regions, but aligned correctly to chrM when the CAST reference was used (Figure 6c).

**Figure 6.**
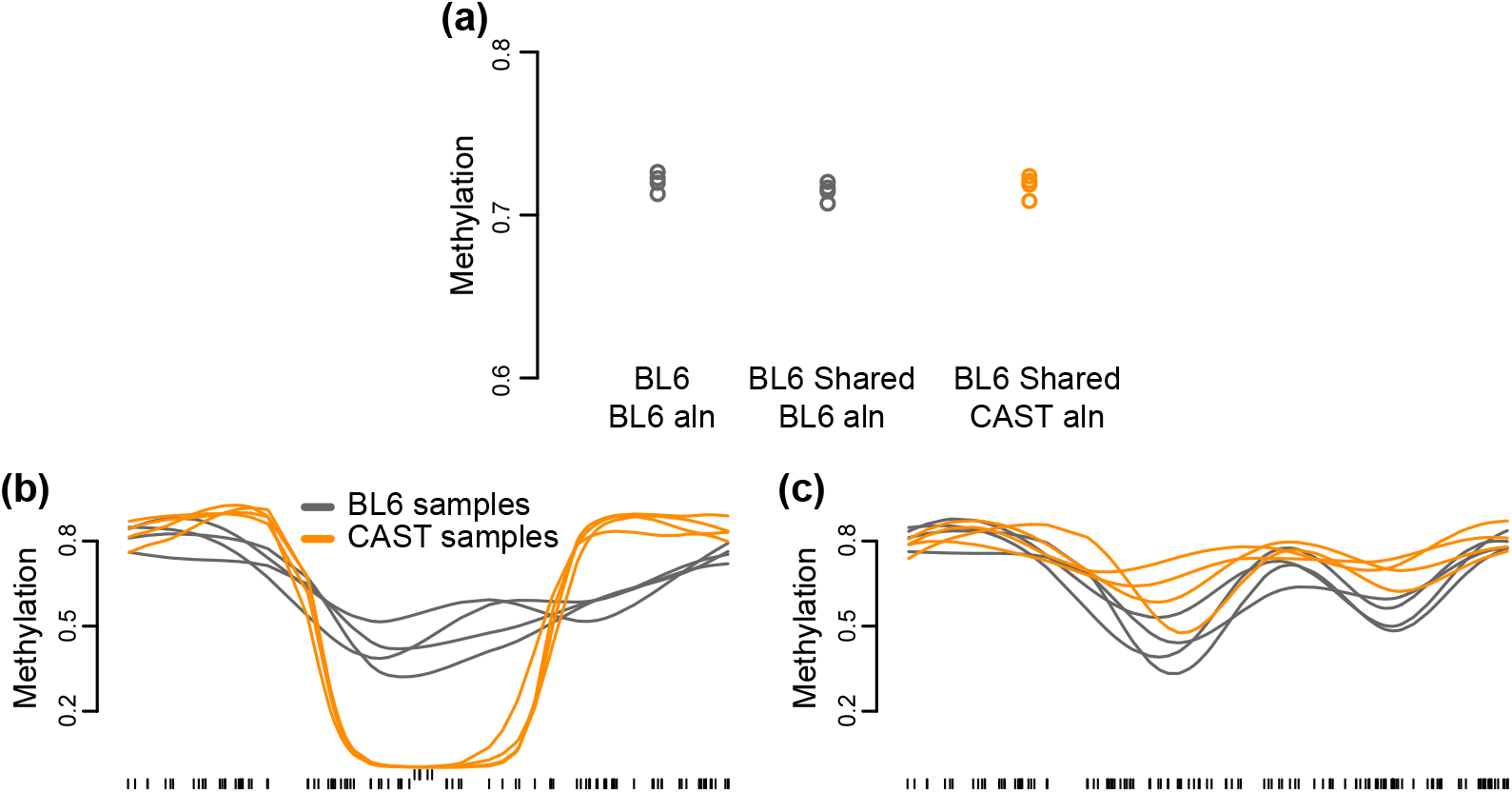
Variation filtering does not completely remove alignment bias. Filtering out strain-unique CpGs from the BL6 reference sufficiently addresses global methylation bias (**a**). However, an example 11kb genomic region, despite containing no CpGs unique to either mouse strains, appears as a DMR under the filtering strategy (**b**) despite no true methylation difference under personal-genome alignment (**c**). This is due to misalignment of reads: upwards tick marks indicate CpGs with average coverage greater than 100 in the filtered analysis.

Overall, when personal reference genomes are unavailable, alternative measures are required to address sites of potential CpG variation. Even then, these measures are likely to result in a loss of power greater than what was observed in our analysis, in which we found fewer regions despite perfect information on strain-specific CpGs. Furthermore, results must be interpreted with caution and careful examination of individual regions, to exclude the possibility of bias from non-CpG alignment errors.

## Discussion

Here we have studied the problem of comparing DNA methylation data between divergent genotypes, at the level of strain differences between inbred mice. We have introduced a method which – through imputation – allows the inclusion of strain-specific CpGs in an analysis of differential methylation. The most powerful version of our method uses strain-specific genomes and handles the analysis of data mapped to different genome coordinates, and we show this results in increased power for detection of strain-associated differentially methylated regions. Our results have clear and important implications for using model organisms to study the interplay between epigenetics (DNA methylation) and genetics (DNA sequence).

In addition to developing our method we have examined the impact of various types of alignment strategies on the ability to quantify strain-associated differential methylation. At the most extreme end, using just a single reference genome results in a massive mapping and quantification bias which completely hides the true signal. We have for the first time precisely quantified this bias by using the strategy of aligning to multiple genomes and observing a reversal of the bias. To detect the bias we recommend routinely examining global methylation across samples. This is easy and effective, but will only detect strong genome-wide mapping bias. Care should be taken; it is well established that global methylation is cell type dependent.

Sometimes, strain-specific genomes are not available and approximations must be made. If a database of known CpG variation is available, methylation calls can be post-alignment filtered for CpGs which vary between genotypes. This results in a loss of power. Our results are best-case results: the available database of CpG variation is perfect. Alternatively, it is possible to use existing tools to genotype samples using bisulfite converted DNA. This was largely unsuccessful, either due to too low coverage (we had 28x) or due to the tool we used.

We have used the term mapping bias throughout this work, because the bias is controlled by the choice of reference genome used for alignment. However, in the case of a C to T transition between the reference genome and the sample genome, the aligner is placing the read correctly, and what fails is our inference based on a combination of the aligned read and the existence of a CpG in the reference genome. Previously, [27] showed that allele specific expression can be affected by mapping bias, whereas [28] showed that eQTL analysis is often unaffected. eQTL analysis is largely unaffected by mapping bias because mapping bias occurs locally around a sequence variant, and gene expression is averaged across a much larger region than is affected by the bias. In contrast, for DNA methylation – like allele-specific expression – the quantity of interest is directly on top of a sequence variant.

The impact of mapping bias depends critically on experimental design. Here we have focused on an analysis where the goal is to compare two distinct strains. A different sce nario would be to compare two groups of mice, with both groups of mice being balanced among different strains; an example could be to compare young to old mice. In such an experiment mapping bias would not cause biased methylation differences, but would rather cause unnecessary between-sample variation within the two groups, with an associated loss of power. The strategy we advocate here would address both situations, removing bias and decreasing variation.

In future work, it will be important to extend this method to the situation where the samples are outbred. Munger et al. [29] explored this in the context of allele-specific expression and found that a personal genome strategy was also effective there. But the task of constructing the personal genome is complicated by the fact that each sample’s genome is a distinct composition of founder haplotypes. This requires the additional step of inferring founder haplotypes across the subject genomes, which incurs additional computational overhead. Depending on the population under study, it may be possible to use auxiliary genetic data to infer this.

In general, we believe that studies in model organisms involving multiple different strains will require the availability of strain specific genomes. This is because such studies are usually undertaken to understand the impact of genotype and therefore involve the comparison of groups of individuals with different (sometimes vastly different) genomes. This is for example the rationale behind the development of the mouse Collaborative Cross [30]. *Arabidopsis thaliana* is frequently used for the same purpose, although plants have extensive non-CpG methylation which we have not considered in our work.

In many human studies different groups of interest are composed of different individuals. We know from genome-wide association studies that individuals are rarely random samples from a background distribution, and such studies are therefore susceptible to being affected by mapping bias. But two different humans are genetically closer than the two mouse strains studied here, and in practice the impact of this bias will depend on the genetic heterogeneity of the samples and the size of the signal of interest.

Our results suggests a tantalizing relationship between regional patterns in DNA methylation and evolution. We observe that many apparent small and large-scale differentially methylated regions reverse when we change the reference genome used for alignment. For a direct reversion to happen, a loss of CpG with an associated methylation level has to be compensated by a nearby CpG gain with a similar methylation level, in such a way that the spatial pattern of DNA methylation is preserved. The prevalence of reversion, and the similar CpG mutation rates between strains, suggests that this might be a general phenomena. Additional work on this hypothesis is clearly required before any conclusions can be drawn.

In conclusion, we have shown that mapping bias can severely affect analysis of bisulfite converted DNA. We have proposed a method which allows for analysis of differential methylation between different genotypes, including genotype-specific CpGs. We show that this increases power but requires the availability of sample specific genomes or at least a database of known CpG variation. Future studies employing bisulfite sequencing need to carefully consider this issue, especially if the goal is comparison between distinct genotypes.

## Material and Methods

### Sample information

Liver samples from two mouse strains (C57BL/6J and CAST/EiJ, 4 mice per strain) were obtained from Jackson Laboratories. All mice were 6-week-old females; additionally, mice of the same strain were littermates.

### DNA extraction and sequencing

Genomic DNA was extracted from liver using the Qiagen DNEasy kit, with an additional RNase incubation step (50 *μ*g/sample, 30 minutes) prior to column application to remove RNA.

WGBS single indexed libraries were generated using the TruSeq DNA LT Sample Preparation Kit (Illumina) according to the manufacturer’s instructions with modifications. *1μ*g gDNA for BL6 samples (1.34*μ*g gDNA for CAST samples due to observed partial DNA degradation) was quantified via Qubit dsDNA BR assay (Invitrogen) and 0.8% agarose gel. 1% Unmethylated lamda DNA (cat#D1521, Promega) was spiked in for monitoring bisulfite conversion efficiency. Samples were fragmented by Covaris S2 sonicator to an average insert size of 350bp (80sec, Duty cycle 10%, Intensity 5, Cycles per burst 200). Size selection was performed using AMPure XP beads and insert sizes of 300-400bp were isolated. Samples were bisulfite converted after size selection using EZ DNA Methylation-Gold Kit (cat#D5005, Zymo) following the manufacturer’s instructions. Amplification was performed following bisulfite conversion using Kapa Hifi Uracil+ (cat#KK282, Kapa Biosystems) polymerase and cycling conditions: 98degC 45s /8cycles: 98degC 15s, 65degC 30s, 72degC 30s / 72degC 1 min.

Final libraries were confirmed via 2100 Bioanalyzer (Agilent) High-Sensitivity DNA assay. Libraries were quantified by qPCR using the Library Quantification Kit for Illumina sequencing platforms (cat#KK4824, Kapa Biosystems), using 7900HT Real Time PCR System (Applied Biosystems).

Libraries were sequenced on an Illumina HiSeq2000 sequencer using 100bp paired-end runs with a control lane.

### Data availability

Data are available under accession number GSE87101 in NCBI GEO. This includes alignments of the samples to both CAST and BL6 genomes, expressed in BL6 coordinates as well as alignment of the samples to the CAST genome, expressed in CAST coordinates.

#### Review link to GEO

https://www.ncbi.nlm.nih.gov/geo/query/acc.cgi?token=qfilaimmzbcpnmf&acc=GSE87101

### Short-read alignment

Alignment and CpG read-level measurement were performed using Bismark version 0.16.1 [5] and Bowtie2 version 2.1.0 [31]. Reference genomes for BL6 and CAST were generated from their corresponding FASTA files (build 37), obtained from UNC Systems Genetics [17]. Sequencing reads were first trimmed using Trim Galore! version 0.3.7 [32] using default options, then aligned with Bismark using options `--bowtie2 --bam`. BAM output files were merged and sorted in preparation for methylation extraction using Samtools version 1.3 [33]. Read-level measurements were obtained using the Bismark methylation extractor, with options `-p --ignore 5 --ignore_r2 5 --ignore_3prime 1 --ignore_3prime_r2 1`. Readmeasurements were modmapped if applicable (see section below), then converted to BSseq objects in R version 3.3.0 using the bsseq package [4]. When creating BSseq objects, forward- and reverse-strand reads were combined for each CpG.

### Coordinate mapping between strains

To facilitate direct comparison of CpGs between strains, we converted genomic coordinates using the “modmap” package from UNC Systems Genetics [18]. This package functions similarly to the liftOver tool [19] on the UCSC Genome Browser, and takes as input two files: 1) the user’s list of genomic coordinates to be converted; 2) a strain-specific MOD file, again obtainable from UNC, describing how to convert coordinates between strains. We used this package to convert a list of all CpG locations in the CAST genome to their corresponding locations in the BL6 genome, and vice versa. CpGs containing negative positions on either strand in the modmap output (indicating the location resided within an insertion or deletion in the other strain) were discarded.

### Genomic mutation rate

The MOD file for CAST lists 20,539,633 bp of single nucleotide variants, 6,633,124 bp of insertions, and 5,279,608 bp of deletions from the BL6 genome to CAST (including autosomes, allosomes, and chrM). The length of the BL6 genome is 2,654,911,517 bp, for an overall genomic mutation rate of (20,539,633 + 6,633,124 + 5,279,608) / 2,654,911,517 = 1.2%.

### Focal methylation analysis

We used the BSmooth pipeline as implemented in the bsseq package from Bioconductor [4], as employed previously [20, 21]. For small DMR analysis, the data was smoothed using BSmooth with the following (default) parameters (ns = 70, h = 1,000, maxGap = 10^8^). Following smoothing, we used t-statistics (cutoff = 4.6, maxGap = 300) to obtain putative differentially methylated regions as described previously, only analyzing CpGs where at least 3 samples in a group had a coverage of at least 2. Significance was assessed using a stringent permutation approach as described previously [21]. Specifically, we used permutations which balanced the two strains (ie., each permutation has 2 mice from each strain in each group); there are 18 such permutations. For each DMR we calculated how many permutations we saw a better null DMR; dividing by the total number of permutations gives us the quantity we call gFWER. To compare DMRs we are searching for DMRs with many large CpG-specific t-statistics; to be precise we say one DMR is better than another if it has a greater number of CpGs as well as a greater total sum of t-statistics across all CpGs in the DMR. By comparing each putative DMR to all null DMRs in each permutation we control for multiple testing and control the familywise error rate, a stringent multiple testing error rate. The interpretation of a gWER of 1/18 for a given DMR is that in 1 out of 18 permutations do we see a bigger permutation DMR *anywhere* in the genome.

### Binned methylation analysis criteria

The mouse genome was divided into 10kb bins. A bin was analyzed for methylation if it had at least 5 covered shared CpGs; 5 covered strain-unique CpGs per strain; and at least a 10% proportion of covered strain-unique CpGs to all covered CpGs in both strains. A CpG was considered covered if it was covered by two or more reads in at least 3 of 4 samples in each strain it is found in. (For example, a shared CpG would need 2+ reads in at least 3 BL6 samples and 3 CAST samples to be considered covered.) 118,553 of 236,632 bins genome-wide fit this criteria.

### Overlap with functional regions

Regions were obtained and defined as described under external data. Given a set of DMRs as well as a class of regions, we compute the odds ratio of enrichment by considering the overlap in CpGs between the two sets of regions, accounting for the fact that not all CpGs where measured in our data. This approach naturally addresses issues of non-uniform distribution of CpGs.

### External data

Genomic intervals for ENCODE/LICR histone [34] and TFBS tracks [35], RefSeq genes [36], and CpG islands [37] were obtained via the UCSC Genome Browser. Histone and TFBS data were generated by the ENCODE Consortium [38] and the Bing Ren laboratory, and are also available at GEO accessions GSE31039 and GSE36027. ENCODE filenames as listed in the UCSC download server are provided in Table 5. Promoter regions of Refseq genes were defined as the 5-kb region flanking a gene’s transcription start site. CpG shores were defined as the 2-kb regions upstream and downstream of a CpG island.

**Table 5.** Filenames for ENCODE data.

| Filename |
| --- |
| wgEncodeLicrHistoneLiverH3k27acMAdult8wksC57bl6StdPk.broadPeak |
| wgEncodeLicrHistoneLiverH3k4me1MAdult8wksC57bl6StdPk.broadPeak |
| wgEncodeLicrHistoneLiverH3k4me3MAdult8wksC57bl6StdPk.broadPeak |
| wgEncodeLicrTfbsLiverCtcfMAdult8wksC57bl6StdPk.broadPeak |
| wgEncodeLicrTfbsLiverPol2MAdult8wksC57bl6StdPk.broadPeak |

## Acknowledgments

We thank Birna Berndsen for assisting with library preparation and sequencing.

## Funding

Research reported in this publication was supported by National Institute of General Medical Sciences of the National Institutes of Health under award number GM118568 (to BL) and the National Cancer Institute of the National Institutes of Health under award number CA054358 (to APF).

## Disclaimer

The content is solely the responsibility of the authors and does not necessarily represent the official views of the National Institutes of Health.

## Competing interests

The authors declare that they have no competing interests.

## Abbreviations

WGBS: whole-genome bisulfite sequencing
DMR: differentially methylated region

